# Nucleotide-specific RNA conformations and dynamics within ribonucleoprotein condensates

**DOI:** 10.1101/2025.02.06.636987

**Authors:** Tong Wang, Qingyue Hu, Scout Fronhofer, Lois Pollack

## Abstract

Ribonucleoprotein (RNP) condensates have distinct physiological and pathological significance, but the structure of RNA within them is not well understood. Using contrast-variation solution X-ray scattering, which discerns only the RNA structures within protein-RNA complexes, alongside ensemble-based structural modeling we characterize the conformational changes of flexible poly-A, poly-U and poly-C single stranded RNA as it interacts with polybasic peptides, eventually forming condensed coacervate mixtures. At high salt concentrations, where macromolecular association is weak, we probe association events that precede the formation of liquid-like droplets. Structural changes occur in RNA that reflect charge screening by the peptides as well as *π* − *π* interactions of the bases with basic residues. At lower salt concentrations, where association is enhanced, poly-A RNA within phase separated RNP mixtures exhibit a broad scattering peak, suggesting subtle ordering. Coarse-grained molecular dynamics simulations are used to elucidate the nucleotide-specific dynamics within RNP condensates. While adenine-rich condensates behave like stable semidilute solutions, uracil-rich RNA condensates appear to be compositionally fluctuating. This approach helps understand how RNA sequence contributes to the molecular grammar of RNA-protein phase separation.

## Introduction

Ribonucleoprotein (RNP) condensates formed through phase separation of RNA and proteins include p-bodies, stress granules, and paraspeckles, and have important functions within cells.^1–5^ RNA modulates the biophysical properties of RNP condensates and can homeostatically drive or buffer their formation. ^6–9^ Although RNA does not require other macromolecular partners for phase separation,^10,11^ physiological condensates often contain both RNA and proteins, and their aberrant condensation and/or subsequent gelation can be pathogenic. ^12–17^

Beyond polymer-solvent incompatibility, RNP phase separation is driven largely by complex coacervation, facilitated by electrostatic attraction between the negatively charged phosphate backbone and positively charged amino acids.^18,19^ Condensate stability is further influenced by hydrogen bonding between nucleotides and pairwise *π*-*π*/cation-*π* interactions of *π* orbitals in aromatic nucleotide bases and basic amino acids (Figure 1A). These so-called “sticker” interactions have been shown to be nucleotide (sequence) specific. Thus, RNA sequence can influence condensate formation by providing a spatially varying (sequence dependent) arrangement of sticker motifs. In addition, single stranded RNA chains of varying sequence have distinctive chain formations, so RNA structure may crucially influence condensate formation.^20,21^ Cellular RNP condensates such as stress granules are enriched in certain RNA sequence and structural features, and different nucleotide/protein sequences and conformations have differing material properties. ^22–24^ Distinct RNA condensate phenotypes may arise such as hollow vesicles and mesh-like arrangements, and mRNA structure has been shown to influence condensate dynamics.^25**?**^ Moreover, RNA clustering and hybridization state can influence condensate morphology and different mRNA constructs form RNP condensates with differing dynamics and viscosities.^26–28^ This behavior reflects the influence of RNA structure on macroscopic RNP condensate properties. We build on this framework, aiming to better understand the interplay between sequence dependence and structural features of highly flexible RNA.

**Figure 1:**
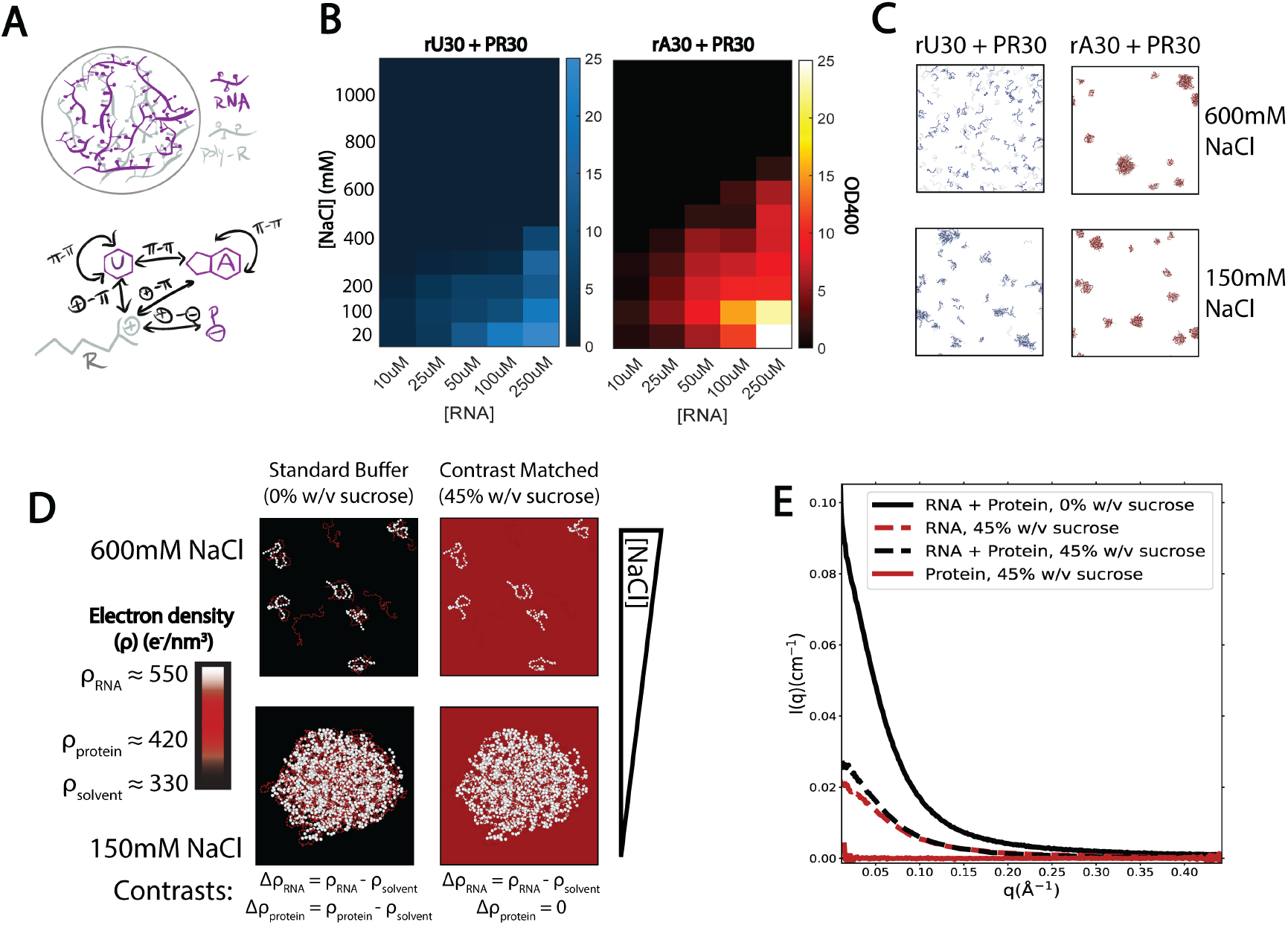
Nucleotide-specific RNA condensate formation propensity in poly-U vs poly-A with PR30 peptide. A) Chemical “sticker” interactions within ribonucleoprotein condensates. *π*−*π* interactions may occur between aromatic nucleotide bases, cation-*π* interactions may occur between nucleotide bases and charged amino acids like arginine (R). Electrostatic interactions may occur between R and the negative phosphate (P) backbone of RNA. In addition to examining the effects of these interactions on RNA structure, we investigate if A vs U interactions have different strengths and lead to different structural morphologies. B) Phase diagrams of rU30 + PR30 and rA30 + PR30 at 100 *µ*M and 250 *µ*M RNA concentrations and a range of NaCl concentrations. Conditions are indicated according to the colorbar, which indicates solution turbidity assayed by measuring absorbance at 400nm. C) Coarse-grained molecular dynamics simulations of rU30 and rA30 with PR30 at 150 mM and 600 mM NaCl concentration, showing the last frame of a 1*µ*s trajectory. RNA is colored blue (rU30) and red (rA30) while PR30 is colored light grey. D) Schematic of the contrast-variation SAXS approach. RNA nucleotides have greater electron density (*ρ*) than peptide amino acids, which normally have sufficient scattering contrast with a standard buffer, whose *ρ* is lower than both. In a contrast-matched solution, *ρ*_*solvent*_ = *ρ*_*poly*−*peptide*_ and only the RNA signal prevails. This study assayed contrast-matched solutions at 600 mM NaCl (near the binodal boundary where subsaturated RNA-peptide complexes form) and 150 mM NaCl (where phase separation into RNP condensates is favorable). E) Solution scattering profiles of an rU30 + PR30 complex in standard buffer vs a contrast matched buffer. The scattering of rU30 and PR30 alone in contrast matched buffer are also shown.

Some work to date has focused on condensates formed with RNA homopolymers, single stranded chains with a monomeric sequence. It has been shown that poly-A, U, and C nucleotides form spherical droplets with poly-arginine proteins characteristic of classical nucleation while poly-G forms a nonuniform fractal-like network more characteristic of a gel.^22^ Past work, studying the conformations of homopolymer RNA chains in solution (in the absence of protein/peptide partners), suggests that they form structures that depend on their sequences, perhaps providing important context for their behavior in condensates. For example, A bases have stronger *π*-*π* stacking propensity than U bases.^29^ Since *π*-*π* contacts are a potential sticker interaction in condensates, this likely contributes to Mg-induced RNA condensates of poly-A forming at lower RNA and salt concentrations than poly-U and having slower dynamics.^30,31^

Given our past structural characterization of homopolymeric single stranded RNA chains, we are well positioned to ask how these chain conformations change upon interaction with coacervation-inducing proteins and how different RNA sequences influence structural behavior in RNP condensates. ^29,32^ While these studies were primarily performed using small-angle X-ray scattering (SAXS), standard SAXS measurements do not easily distinguish signals from different macromolecular components. We therefore turn to contrast-variation (CV) SAXS. With roots in experimental condensed matter physics, solution X-ray and neutron scattering have probed structural features in systems of densely interacting polymers such as amorphous solids, liquid crystals, and segregated mixtures. ^33^ CV-SAXS offers the added ability to highlight the conformation of one component by ‘blanking out’ the other components in a multispecies mixture with different electronic densities (such as an RNA-protein mixture). It has previously been applied to examine nucleic acid conformations when complexed with RNA-binding proteins, nucleosomes, and viral capsids.^34–37^

Here we employ CV-SAXS to measure the conformations of poly-U, poly-C, and poly-A RNA of 30 nucleotide lengths in complex with a pathogenically relevant poly(proline-arginine) (PR) repeat peptide. ^22,38–40^ These molecules will subsequently be referred to as rU30, rC30 and rA30. CV-SAXS reports the structural features of just the RNA within the mixture. Using this method, we characterize changes in rU30, rC30, and rA30 chain conformation at high NaCl concentration, where coacervation begins to occur. Because our past studies focused on the conformations of rU30 and rA30 (and not yet on rC30, though work is in progress), we focus our analysis on changes beginning with known conformational ensembles. At closer to physiological NaCl levels where RNP droplets form, we observe a more drastic transition with a diffuse scattering peak in the SAXS profile that is unique to poly-A. Through coarse-grained MD simulations using the Mpipi model, ^41^ we further generate models of phase-separated RNA and show that poly-A forms stable condensates, while poly-U and poly-C condensates are compositionally fluctuating. The critical importance of RNA within biomolecular condensates in physiology and disease, coupled with the growing amount of RNA-centric studies in the field, highlights the need for studies that expand the fundamental framework of RNA phase transitions.

## Materials and methods

### RNA-PR30 phase separation

To induce phase separation, 250*µ*M rU30, rA30, or rC30 RNA was mixed with PR30 at 2:1 PR30:RNA stoichiometric ratio to ensure that all (−) charges on RNA have at least one (+) partner. NaCl concentrations of 150 mM and 600 mM were tested in this study. RNA was added last to solutions containing the prescribed amounts of of NaCl and PR30. Mixing was performed by pipetting half the sample volume up and down 20-25 times. The occurrence of phase separation was assessed by monitoring solution turbidity through absorbance measurements at 400nm using UV-Vis spectroscopy. Phase diagrams, displayed on axes of {[RNA],[NaCl]}, were measured by reconstituting RNA-PR30 mixtures at 20 mM NaCl and titrating, adding a stock of 5 M NaCl, in 100 mM increments up to 1 M NaCl. A new RNA-PR30 mixture was prepared halfway through this titration so that the concentration was never reduced by more than 20%. To ensure that this method of titrating NaCl was accurate, one phase diagram, measured as described above, was compared with another one, where each condition was assessed as a separate reaction. For this latter phase diagram, reactions were set up in parallel on a 96-well plate using a multichannel pipette. A SpectraMax M2 (Molecular Devices, San Jose, CA, USA) plate reader was used to measure absorbance at 400 nm for each reaction simultaneously. Both measurement protocols yielded similar phase boundaries and RNA/NaCl saturation concentrations (Figure S1). Since coacervation was reversible and the titration method uses significantly less sample at the high RNA concentrations probed, we used this method to build phase diagrams. Turbid mixtures were confirmed to contain phase separated droplets through brightfield microscopy on a Zeiss Axiovert 135, imaging through a UMPlanFl 20x/0.50W objective lens and recording images using a 1.31 MPix 1/1.8” CMOS camera (UI-3240CP Rev. 2, Edmund Optics). Droplets were monitored periodically for 3-60 minutes after mixing to observe coarsening and fusion.

### Peptide match point determination and verification

With equation (S.4) in mind, we estimated the match point of PR30, the sucrose concentration that raises the buffer electron density until it equals that of the peptide, by measuring I(0) of 10mg/mL peptide samples in 150 mM NaCl, 1mM MOPS (pH 7.07), 20*µ*M EDTA buffers. Measurements were performed with ascending amounts of sucrose - 0, 10, 20 %w/v (Figure S2A). We then plotted 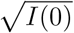 against buffer electron density (*ρ*_*buffer*_) and linearly extrapolated the *ρ*_*buffer*_ at which 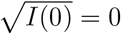 (Figure S2B). *ρ*_*buffer*_ was then converted to % w/v sucrose using equation (S.5). To obtain values for I(0), SAXS experiments were performed on a BioXolver laboratory X-ray source (Xenocs, Grenoble France), averaging 12 × 180 second exposures for each sample at 100% X-ray intensity. The images were azimuthally integrated to plot the scattering intensity profile I(q). The y-intercept, I(0) was determined from a Guinier fit for *qR*_*g*_ < 1.3:

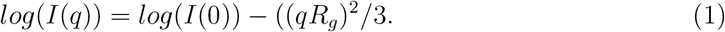

where *R*_*g*_ is the radius of gyration.

To verify the match point, BioXolver SAXS measurements on PR30 were performed at 45% and 50% w/v sucrose, near the extrapolated *c*_*wv,sucrose*_, and the PR30 vs buffer scattering profiles were examined for intensity differences (Figure S2C,D). Samples were prepared at the correct concentration of sucrose by pipetting in a high sucrose concentration stock (70% w/v buffer). 45% sucrose was observed to result in PR30 scattering that was more similar to the respective buffer scattering (Figure S2C,D), so this was determined to be the match point of PR30. The same measurements were performed at both 150 mM and 400 mM NaCl to determine whether the match point is salt dependent (Figure S2C,D).

### Contrast-variation solution X-ray scattering

Prior to CV-SAXS measurements, dialysis was performed to ensure that the same sucrose concentration was present in both the RNA and peptide samples. A target buffer (the experimental buffer with 45% w/v sucrose) was prepared before dialysis. 100 *µ*L of each RNA and peptide sample was premixed with 100 *µ*L target buffer in a pre-washed 350 *µ*L dialysis button (HR3-332, Hampton Research) to reduce the dialysis time. The buttons were sealed carefully with a pre-wet (in the target buffer) 3.5k MWCO SnakeSkin dialysis tubing (Thermo Scientific) and placed in separate floating racks (1 rack for rU30 and rA30, and 1 rack for rC30 and PR30). Each rack was placed in a 150 mL beaker containing 120 mL target buffer at 4°C vertically under the liquid surface to maximize the dialysis efficiency. The buffer in the two beakers was then mixed using a magnetic stir bar for 20 hours at 4°C at 400 rpm. This dialysis duration was required for accurate buffer matching in SAXS measurements. To collect the sample after dialysis, the dialysis buttons were removed from the buffer, and a plastic syringe with a needle was used to pierce through the dialysis membrane and transfer the sample into different Eppendorf tubes. Before the measurements, the RNA and peptide concentrations were adjusted to 250 *µ*M and 15 mg/mL respectively.

CV-SAXS experiments were performed at the ID7A1 BioSAXS/high pressure SAXS beamline at the Cornell High Energy Synchrotron Source. ^42^ As the addition of sucrose decreases the signal strength by reducing contrast, beam attenuation was varied to maximize signal without radiation damage. Data were collected through 100 × 1 second X-ray exposures, monitoring radiation damage by observing stability of traces over multiple exposures. Beam centering and azimuthal integration of scattering images to obtain 1D I(q) scattering profiles were performed using BioXTAS RAW.^43^ Data are shown on Kratky (I(q) scaled by *q*^2^) and normalized Kratky (q scaled by *R*_*g*_ and I(q) scaled by (*qR*_*g*_)^2^/I(0) to normalize out size information). More information on SAXS data acquisition is provided in Supplementary Table 1.^44^

To ensure thorough cleaning of the sample cell, particularly for phase separated samples with added sucrose, two consecutive cleaning cycles were performed between each measurement. For each cycle, we flowed a solution containing 10% methanol, 2% hellmanex for 10 seconds, pure methanol for 10 seconds, and Milli-Q water for 30 seconds. Cleaning was verified periodically by acquiring water or buffer background scattering profiles and ensuring that the signals did not change. Any increase in background would indicate sample cell fouling. The quality of background subtraction of sucrose containing samples was monitored throughout the experiment by assessing alignment of buffer vs buffer + macromolecule traces at high q (q~0.3*Å*), looking for lack of unphysical buffer crossover events where the buffer scattering exceeds the buffer + macromolecule scattering (Figure S3A-C). Buffer matching quality was also assessed by measuring RNA-alone SAXS profiles at 600 mM NaCl and comparing them to corresponding SAXS profiles collected with no sucrose (Figure S3D,E).

### Disordered RNA chain builder and ensemble optimization method (EOM)

To identify structural ensembles that recapitulate our CV-SAXS data, we used dinucleotides as the fundamental building block to construct RNA chains. The dinucleotides are parameterized by their {*α, β, γ, ζ, ϵ*} torsional angles, {*δ, δ*(i − 1)} sugar puckers, and {*χ, χ*(i − 1)} nucleotide orientation angles. RNA dinucleotides arrange into only a finite number of angle/sugar pucker combinations due to steric biases, and these RNA ‘suites’, or angle sets, have been defined previously.^45^ Only 34 sets of torsional angles that are common to singlestranded RNA are employed, thus the overall parameter space consists of 34 suites of 11 variables for every dinucleotide unit. 30mer RNA chains are built using 29 consecutive dinucleotide units, assigning a suite to each unit. Suites are assigned initially using a Gibbs sampler, disallowing steric violations. The method is described in detail by Plumridge et al. ^46^ A starting pool of 2000 30-mer poly-U or poly-A was generated (29 suites per conformer), where the weights of the RNA suites are approximately equal across the ensemble and the same set of suite weights was initially used for all cases. CRYSOL v2.0 was used to compute the theoretical scattering profile of all conformers, with a maximum order of harmonics of 15, Fibonacci grid order of 18, 0.3 *Å*^−1^ maximum scattering angle, and 61 calculated data points.^47^ GAJOE v1.3 was then used to determine the subset of the ensemble that agrees with the experimental data, by running 1000 generations with 50 ensembles, 20 curves and 10 mutations per ensemble, over 100 iterations.^48,49^ For the algorithm, repeat selections were allowed but I(q) curve offsets were not.

At each step, M structures are selected from the full starting pool and the linear combination of SAXS profiles {*I*_*k*_(*q*); *kϵ*[1, *M*]} is computed:

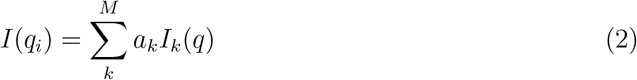

The fit to the scattering data was assessed by computing the *χ*^2^ difference between theoretical and experimental SAXS profiles.

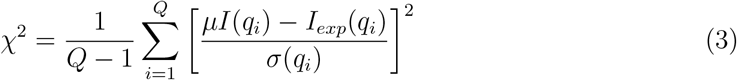

Here, Q is the number of points sampled in q-space, *I*_*exp*_(*q*_*i*_) is the experimental SAXS data with uncertainty *σ, µ* is a scaling factor, and *I*(*q*_*i*_) is given by equation (2).

The process was iterated 9-10 times, each iteration reshuffling the dinucleotide suite weights (w) such that the discrepancy between observed suite frequencies (*h*_*ens*_) and their expected values is minimized.

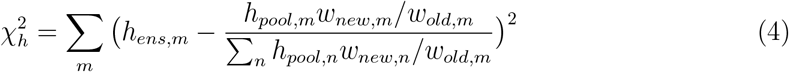

Where *h*_*pool*_ is the frequency of suites in the full pool.

The reduced chi square was computed to account for the global fit of all N=50 ensembles.

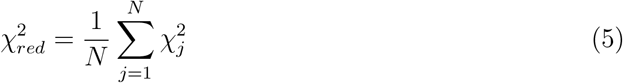

As a secondary metric to assess convergence of the iterative refinement procedure, changes in the distributions of RNA suite weights (W) between iteration i and i+1 were quantified using the Jensen-Shannon divergence: ^50^

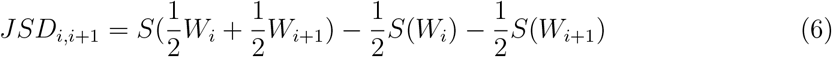

This allows us to assesses how close RNA suite weight distributions are between consecutive iterations. S(W) is the Shannon entropy for the discrete weights probability distribution W.

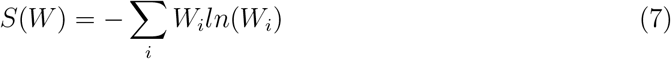

For validation, two independent rounds of EOM were performed, yielding comparable results.

### Structural order parameters

To visualize ensemble-level structural characteristics in the EOM RNA pools we utilized the orientation correlation function (OCF), which signals the presence or absence of periodic order and helicity along a polymer chain.^51^ The OCF was computed for each RNA chain as:

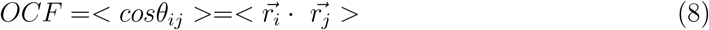

Where 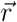 are the normalized bond vectors between each pair of backbone phosphorus atoms, and the dot product is computed for every pair of bond vectors {*i, j*}. OCFs were plotted as a function of the distance between linkages |*i − j*| for all {*i, j*} pairs. For a metric of approximate polymer stiffness and degree of structure, correlation lengths (*l*_*OCF*_) were computed by summing across OCFs in equation (8).

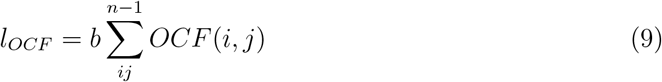

where b is the bond length between sampled coordinates.

Moreover, to estimate the prevalence of intramolecular *π* − *π* interactions within RNA chains, the number of base stacking events was computed for each structure. High base stacking, particularly among many adjacent bases in a row, can indicate nucleotide order along the chain. Purine or pyrimidine bases were indexed by sampling the coordinates of non-hydrogen atoms within the aromatic nucleotide rings. These coordinates were used to fit a plane to represent each nucleotide, using multilinear regression. The normal vectors of each nucleotide plane were then compared, and pairs of bases were considered stacked when their normal vectors were ≤5 *Å* apart and approximately colinear, with ≤45°angular separation. This nonzero angular separation upper limit accounts for nucleotide twist with backbone periodicity. A visualization of base stacking determination is shown in Figure S6. Both OCF and % base stacking were computed as weighted means across all structures in the ensembles, weighted by how much each structure was chosen in the EOM algorithm. All structural descriptors were computed using in-house software written in MATLAB R2021a (Mathworks, Natick, MA, USA).

### Molecular dynamics simulations

Residue-level coarse-grained MD simulations (one bead per nucleotide or amino acid) were performed in Large-Scale Atomic/Molecular Massively Parallel Simulator (LAMMPS, 23 Jun 2022 - Update 3) using a Langevin integrator with a relaxation time of 100 ps for simulations studying large condensates and 5 ps for simulations emulating phase separation propensity with salt. ^52,53^ A step time of 10 fs was used, and simulations were run for 1E8 steps (1 *µ*s) at 298K temperature. A harmonic bond potential was defined between every pair of consecutive residues:

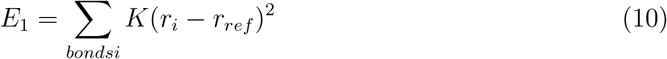

with spring constant K = 9.6 *kJ*/(*molÅ*^2^) and bond distances *r*_*ref*_ = 5.00*Å* for nucleic acids and *r*_*ref*_ = 3.81*Å* for amino acids.

Electrostatic interactions were considered through Debye-Huckel theory: ^54^

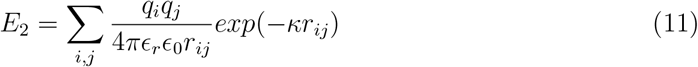

Residue masses, charges, and pair potentials for RNA nucleotides and PR amino acids were defined as specified in the Mpipi model, developed by Joseph et al. ^41^ In this model, a Wang-Frenkel potential is introduced to consider *π* − *π* interactions, which have been shown to be prominent in biomolecular condensation, including that involving RNA.^55^

To emulate experimental conditions performed under ambient temperature and two salt concentrations, the temperature was fixed at 298 K and 600 mM and 150 mM monovalent ion concentrations were specified implicitly through the Debye-Huckel screening length. Screening lengths of 0.253 and 0.126 *Å*^−1^, respectively were employed, calculated from the original formalism: ^54^

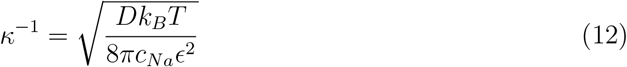

which at standard conditions, with *k*_*B*_ = 1.38E-23 J/K, T = 298K, *ϵ* = 4.77E-10 *cm*^3/2^*g*^1/2^*s*^−1^ ion charge, and ion concentration *c*_*Na*_ in mol/L units, simplifies to:

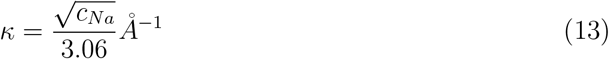

RNA concentrations were set to 250 *µ*M with equimolar PR30 concentrations in a 792.7 *Å* x 792.7 *Å* x 792.7 *Å* box, with the initial condition being RNA and peptide chains starting equally spaced in a cubic lattice. For subsequent analyses of the simulation results, we focused only on RNA coordinates, computationally analogous to blanking the protein signal in the CV-SAXS measurements.

Clusters of RNA chains were identified using ‘Cluster analysis’ in OVITO v3.10.3 with a cutoff distance of 7.5 *Å* (1.5x the RNA bond length).^56^ Cluster coordinates were unwrapped to undo discontinuities at periodic boundaries on a frame by frame basis, so that the time evolution of continuous clusters could be analyzed. All other post-processing of MD simulations was written in house in MATLAB v2023b (Mathworks, Natick, USA). OVITO was also used to visualize MD simulations.^56^

### Condensate formation and exchange rate quantification

Growth and number fluctuation of RNA condensates was considered by counting the number of RNA strands in each cluster for each frame of the simulation, as well as the number of clusters per frame.

To consider the exchange of RNA strands into and out of initial RNA-peptide assemblies and later stage condensates, we considered the set of *N*_*RNA*_ RNA strands within each frame of the simulation. An adjacency matrix *A*_*RNA,f*_ was defined for each frame f, whose elements are *A*_*i,j*_; (*i, j*)*ϵ*[1, *N*_*RNA,f*_]. *A*_*i,j*_ = 1 if *RNA*_*i*_ and *RNA*_*j*_ are in the same cluster, and *A*_*i,j*_ = 0 otherwise. A migration matrix *M*_*RNA,f*_ was then computed by taking the difference of *A*_*RNA,f*_ between adjacent frames:

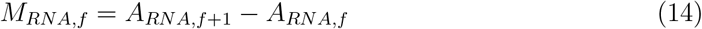

If migration of an RNA strand from one cluster to another occurred, or dissociation of an RNA strand from a large cluster into free solution/association of a single RNA strand from free solution, a nonzero value would manifest in *M*_*RNA,f*(*i,j*)_. Specifically, a positive value indicates an RNA arrival event and a negative value indicates a departure. To prevent overcounting, we discluded all but one arrival/departure event from each individual row or column of M.

## Results

### RNA condensate formation propensity is nucleotide specific

Just as segregation effects in Flory phase separation are modulated by polymer concentration and temperature, coacervation-induced phase separation can be varied by adjusting monovalent ion (NaCl) concentration. At higher concentrations, NaCl competes with contacts between condensed polyelectrolytes and phase separation is less favorable, although polymers may still interact.^18^ Before measuring the structure of RNA, we searched for solution conditions where phase separation occurs by making phase diagrams.

Working in {[RNA],[NaCl]} phase space, the binodal boundary of PR30-induced coacervation with rA30 occurs at a higher NaCl concentration than with rU30 and rC30 (Figure 1B,S7A,B). Specifically, at the 250*µ*M RNA concentration required to measure a robust scattering signal, the turbidity of the solution containing rA30 vanished at 600-700 mM NaCl, approximately 200-300 mM NaCl higher in concentration than the solution that instead contained rU30 or rC30. This provides a first indication of sequence-dependent differences in RNA coacervation: coacervates containing rA30 are more resistant to cation denaturation than those containing rU30 and rC30. As a first test of this distinct behavior, RNA-peptide association was simulated at 600 mM and 150 mM NaCl (see Materials and Methods section **Molecular Dynamics Simulations**), where condensates were observed in both salt conditions for rA30 but not for rU30 (Figure 1C). Simulations are further analyzed later in this study.

The addition of 45% w/v sucrose in buffer solutions, required for CV-SAXS experiments, did not disrupt droplet formation in rU30, rA30, and rC30 complexed with PR30 peptide at 150 mM NaCl and 250 *µ*M RNA concentration (Figure S8). As in the absence of sucrose, these droplets were spherical, coarsened over time, and fused upon contact. At 400 mM NaCl, droplets were observed only with rA30 but not rU30 or rC30, suggesting that the binodal boundary shifts slightly downward in the presence of sucrose. Although the addition of sucrose slightly varies the onset of the phase separation, it does not seriously compromise the droplet morphologies and liquid-like behavior.

Based on the phase diagrams, we selected NaCl concentrations of 600 and 150 mM for our structural studies. The former concentration probes the system above the phase boundaries of rA, rU, and rC condensates at our selected RNA concentration, and allows for the observation of pre-condensation RNA-protein interactions. The latter salt concentration, in addition to being the physiological monovalent ion concentration, provides a condition where complete condensation is favorable both without sucrose and with the addition of 45% w/v sucrose.

### Changes in RNA polymeric state upon peptide association in subsaturated complexes

To explore whether the different structural features of polyrA vs polyrU and polyrC could contribute to the salt dependence described in the previous section, we performed CV-SAXS on multiphasic mixtures of RNA and PR30 (Figure 1D). The first critical step in CV-SAXS studies is to identify the match point, in this case the sucrose concentration where the scattering contribution from the peptide vanishes. We performed a sucrose series and linearly extrapolated a plot of 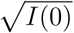 vs *c*_*wv,sucrose*_, as described in Methods section **Peptide match point determination and verification**, and determined that the match occurs near 45 %w/v sucrose. Scattering measurements of PR30 performed at 45% w/v and 50% w/v sucrose confirmed this extrapolation. The PR30 scattering signal was better matched at 45% w/v sucrose (Figure S2C,D) so we selected this concentration for all studies. This agreement held at both 150 mM and 400 mM NaCl, suggesting that the match point is insensitive to increases in NaCl in the concentration regime used in this study. At the match point where the PR30 signal is indistinguishable from the background, the RNA scattering signal prevails (Figure 1E). As a control, we measured the scattering profiles of RNA-alone under that match condition and compared them to those collected in 0% w/v sucrose solution (Figure S3A-E). These results confirm that the addition of 45% w/v sucrose does not appreciably influence the inherent conformations of disordered single-stranded RNA.

At the higher 600 mM NaCl concentration, where full coacervation does not occur, RNA-peptide association occurs without excessive multimerization. This is assessed from the 0% sucrose scattering profiles that show only modest I(0) increases, and because the 45% sucrose scattering profile (reflecting only the RNA) suggest that the RNA remains mostly monomeric (Figure S4, S5). We can thus apply our established ensemble-modeling methods to derive RNA conformations as it interacts with the peptide. This approach not only enables the interpretation of differences in RNA chain compaction with the addition of peptide, it also provides insight into the earliest stages of the RNA-peptide association that precedes complete coacervation. At 600 mM NaCl, the single stranded RNA chains behave like a dilute polymer in a good solvent. This is confirmed by Gaussian nature of the form factor at low q, suggestive of solution monodispersity, as well as the value of the fitted Flory parameters that exceed 0.5 (Figure 2A, S7C, Supporting Methods section **Observables from SAXS**).^57,58^ With the addition of PR30 peptide, the scattering profile (form factor) of RNA changes without a considerable increase in I(0), suggesting a conformational change as opposed association with other RNAs. Association with the protein can be assessed only from the no sucrose scattering profile. (Figure 2A, S5, S4). For rU30, the *R*_*g*_ increases by 8.5% from 23.25 ± 0.24 *Å* to 25.22 ± 0.24 *Å* and *ν* changes from 0.551 ± 0.003 to 0.492 ± 0.002. If using the classical Flory interpretation, ^57^ rU30 crosses the theta point into the poor solvent regime upon the addition of peptide, where intra-polymer association is predicted to be favorable, however the actual trend in RNA chain conformations may be more nuanced than this (Figure 2A,B). For rA30, *R*_*g*_ increases by 11.4% from 21.41 ± 0.21 *Å* to 23.85 ± 0.25 *Å* and *ν* changes from 0.536 ± 0.004 to 0.514 ± 0.003 (Figure 2A,B). Overall, when interacting with peptide, both poly-A and poly-U subtly extend, their profiles display a lower *ν*, or a greater Porod exponent. The same trend is seen for rC30 which undergoes an increase in *ν* from 0.507 to 0.559 and an increase in *R*_*g*_ from 23.53 ± 0.20 *Å* to 24.73 ± 0.24 *Å* (Figure S7C,D).

**Figure 2:**
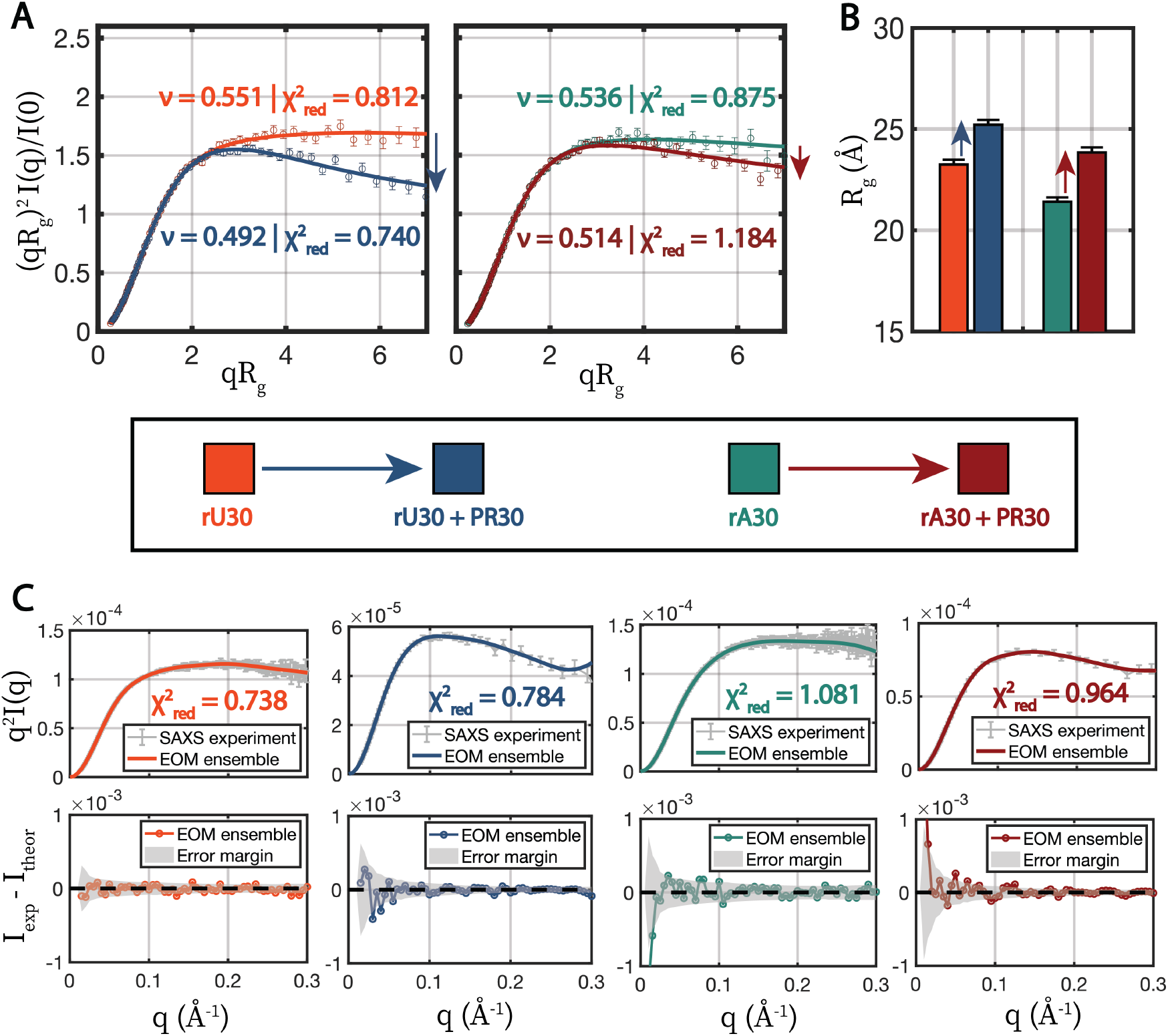
SAXS on rU30 and rA30 RNA + PR30 in contrast-matched solutions at 600 mM NaCl in sub-saturated complexes. Conditions are indicated by color, according to the legend in the center. A) Scattering plots showing changes in conformational states with the addition of PR30. Data are plotted on normalized Kratky axes (see Materials and Methods section **Contrast-variation solution X-ray scattering**), which emphasize high q scattering changes and normalize out size information, focusing on changes in shape. CV-SAXS data are shown as open circles with errorbars and MFF fits are shown as solid lines. Fitted Flory values (*ν*) are listed next to their corresponding datasets, with an accompanying reduced *χ*^2^ from the fit. Errorbars show SAXS measurement errors. B) Guinier-derived radii of gyration of RNA with and without PR30. Errorbars show errors in linear Guinier fits. C) Fits of structural pools determined through optimization with the experimental data on the four conditions assayed. CV-SAXS data are shown in grey on Kratky axes, which emphasize subtle changes at high q, and the total theoretical scattering of each structural ensemble is shown as solid colored lines matching their condition, with accompanying *χ*^2^ values. Errorbars reflect SAXS measurement errors. Residuals between theoretical vs experimental scattering are shown in the bottom plots, with the experimental error margin highlighted in grey.

### RNA backbone and nucleotides undergo conformational changes upon PR30 interaction

The scattering profiles, described above, clearly indicate changes in RNA conformation following association with the peptide. Changes in *R*_*g*_ and *ν* refer to bulk, average properties and are limited in providing more detailed information about RNA conformations, though they are consistent with other approaches. Specifically we speculated how decreases in *ν*, which are often indicative of chain compaction, ^58,59^ are accompanied by increases in *R*_*g*_. To gain more detailed insight into RNA chain conformation using our CV-SAXS results at 600 mM NaCl, we performed an iterative ensemble optimization approach that generates pools of all-atom RNA structures whose theoretical scattering agrees with our experimental measurements (Methods section **Disordered RNA chain builder and ensemble optimization method (EOM)**). Most importantly, this analysis is performed on the RNA both in the absence and presence of the peptide, to discern any peptide-induced changes in the RNA conformational ensemble. This method yields conformational ensembles that agree with our scattering data both in the absence and presence of PR30 (Figure 2C, S7E,F). Across iterations, the 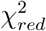 either decreased to convergence or did not markedly fluctuate and Jensen-Shannon divergences generally converged or settled to low values, signifying that the ensembles were no longer changing (Figure S9). Structural ensembles are shown in 2D {*Rg, R*_*EE*_} space in Figure S10, and recapitulate the global structural measurements reported above: similar increases in *R*_*g*_ are seen in the RNA ensemble in the presence of PR30. Interestingly, a new area of the {*Rg, R*_*EE*_} diagram becomes populated when the PR30+ ensemble is considered, signifying larger, more extended RNA chains. This effect is more pronounced for rA30 and rC30 (Figure S10).

As in past work, we mined the predicted ensembles, using ensemble-level orientation correlation functions (OCF), which provide more insight into overall chain configurations and repeated features, ^21,29,51^ as well as an analysis of base stacking. In 600 mM NaCl in the absence of peptide, the OCF of rU30 exhibits an exponential decay characteristic of a random chain without many repeated features (Figure 3A). Under the same conditions, the OCF of rA30 exhibits oscillatory behavior suggesting helicity with a periodicity of 8-10 bases (Figure 3B). rA30 also displays more base stacking than rU30; our analysis reports 70.23 ± 2.92 % adenine bases stacked compared to 37.35 ± 3.89 % uracil bases stacked (Figure 3C,D). This analysis agrees with previous characterizations of these two homopolymeric sequences. In that work, poly-A chains displayed prevalent single-stranded A-form helical conformations stemming from *π* − *π* stacking between adjacent bases.^29^ A more recent study of single stranded RNA conformations exploited graph theoretic class averaging to determine real space RNA conformers in each condition - full structural ensembles are visualized in Figure S11.^21,60^ An examination of the representative structures shows that the adenine bases in rA30 are visually more ordered along the chain than uracil bases in rU30, orienting inward in parallel stacking arrangements that reflect (or possibly cause) backbone ordering (Figure 3E,F). Conversely, the bases in rU30 are more randomly oriented, and the backbone exhibits less ordered helicity (Figure 3E). These features are completely consistent with the more disordered OCF and lower base stacking in rU30.

**Figure 3:**
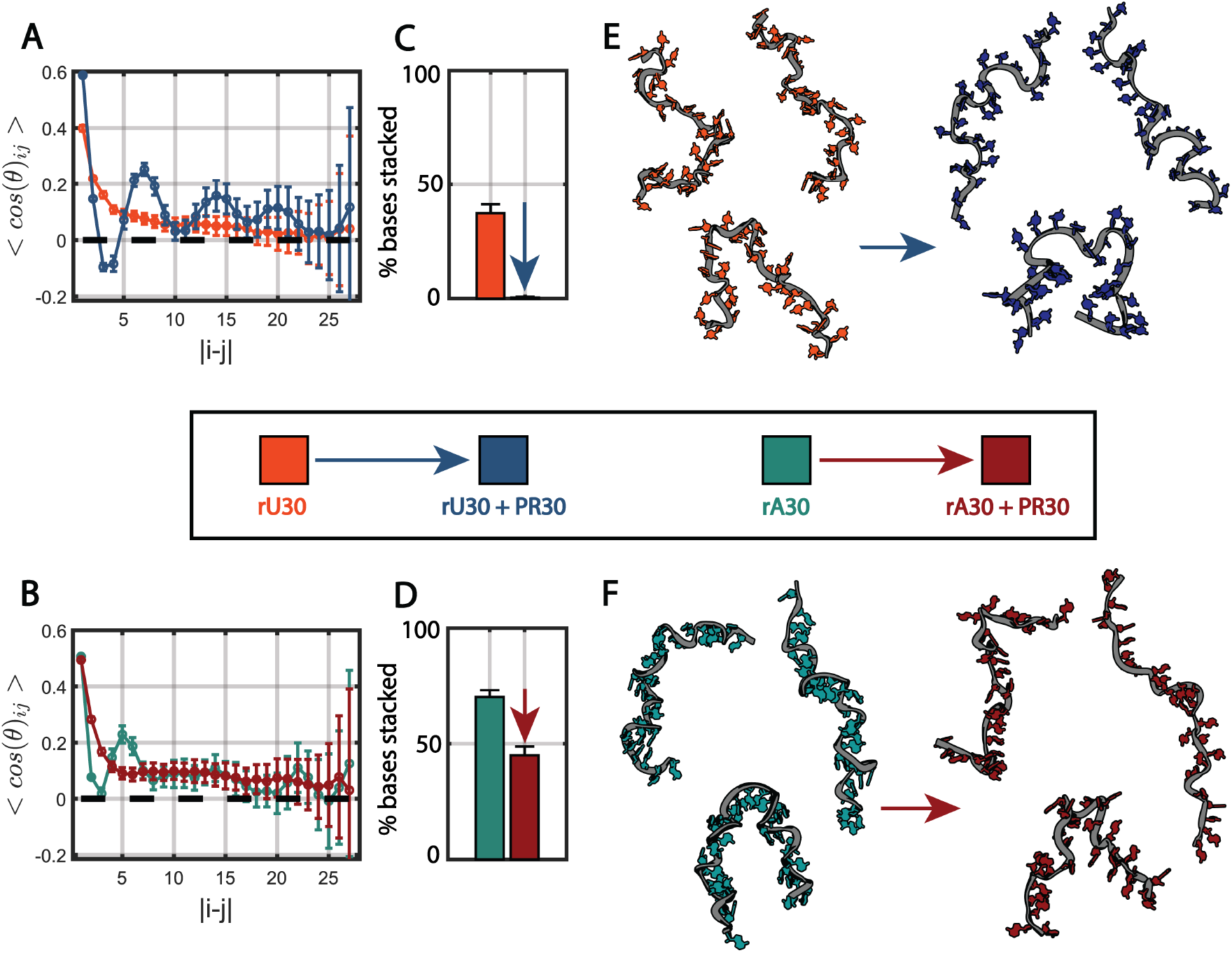
rU30 and rA30 conformational changes with PR30 at 600 mM NaCl in sub-saturated complexes. Conditions are indicated according to the color in the legend in the center. A-B) Orientation correlation function of rU30 and rA30 chains alone and in the presence of PR30. Errorbars show variance across the structural ensemble. C-D) Mean percent of bases within rU30 and rA30 that are stacked without vs with PR30. Errorbars show standard deviations. E-F) Representative all-atom conformers of rU30 and rA30 alone vs with PR30. Nucleotide bases are colored consistent with the legend in the center. Phosphate backbones are colored grey and shown as ribbons for simplified viewing. Each conformer is a centroid structure after spectrally clustering the structural ensembles (see Supplemental Methods section **Conformer visualization**).

Dramatic changes are seen when the same analysis is applied to the contrast matched RNA scattering profiles, following the addition of PR30. When peptide is added rU30 base stacking is completely abolished, with 0.39 ± 0.41 % of bases, virtually none, stacked (Figure 3C). rA30 base stacking is also decreased to 45.01 ± 3.80 % (Figure 3D). The OCF of rU30 transitions now displays distinct periodicity while the OCF oscillation in rA30 is no longer as pronounced, suggesting that rA30 loses its periodic helicity while rU30 gains strong helical features. Across its entire N = 312 structure ensemble these features in rU30 average into a strongly oscillatory OCF with a periodicity of 10-12 bases (Figure 3A, S9). The *l*_*OCF*_ of rU30 goes from 11.98 ± 0.07 *Å* to 18.66 ± 0.07 *Å* with the addition of PR30 and the *l*_*OCF*_ of rA30 goes from 15.00 ± 0.07 *Å* to 17.52 ± 0.08 *Å*, supporting the notion that rU30 chains gain more order when interacting with PR30 while rA30 chains retain a similar amount, even with reduced helicity.

Upon closer examination of the RNA dinucleotide suites selected for the fitting of the rU30 ensemble with PR30 present, over 24% of the dinucleotides have their torsional angles in a conformation that produces an exotic helix that is neither A-form nor B-form (Figure S9).^29,45^ In this dinucleotide arrangement, the uracil bases are also oriented outward relative to the phosphate backbone. When visualizing the all-atom conformations, the bases, for both rU30 and rA30, adopt an ordered arrangement facing away from the backbone (Figure 3E,F). In terms of the RNA backbone twists, rU30 gains a more ordered helical arrangement that compliments its base organization, while rA30 loses some of its characteristic helicity, seemingly because intramolecular base stacking is compromised. Trends with rC30 are similar - base stacking decreases markedly from 61.98 ± 3.43 % of bases stacked to 13.70 ± 2.54 % with PR30. This is reflected in the decreased OCF peak and loss of helical features with bases flipping outward in the real space conformers (Figure S12A-D).

Overall, RNA strands adopt unusual conformations when interacting with polybasic PR30 - the bases flip outward and intramolecular base stacking is reduced. These conformations may result from novel *π* − *π* and cation-*π* interactions between nucleotide bases and arginine in PR30. This structural description enhances the *R*_*g*_ and *ν* changes described in the previous section.

### RNA ordering uniquely accompanies the poly-A phase transition at physiological NaCl

Having examined conformational changes of individual (non-associating) RNA molecules during initial stages of PR30 interaction at 600 mM NaCl, we turn now to the behavior of phase-separated assemblies. At a lower salt concentration of 150 mM NaCl, RNA + PR30 samples exhibit greater turbidity and form liquid-like droplets seconds after mixing. Droplet formation occurs in solutions containing 0% or 45% sucrose (Figure S8). In this region of phase space, far from the binodal in both rA30 and rU30, RNA is expected to form multimeric condensates instead of lower stoichiometry complexes like those observed at 600 mM NaCl.

Our CV-SAXS results on multiphase RNA-peptide samples at 150 mM NaCl show large upturns in scattering at low-q, reporting association of RNA into larger structures under these conditions (Figure 4A). As the sample is no longer monodisperse, the Guinier, polymer fit, and EOM analyses performed at 600 mM NaCl do not provide meaningful interpretations of the data at 150 mM NaCl.

**Figure 4:**
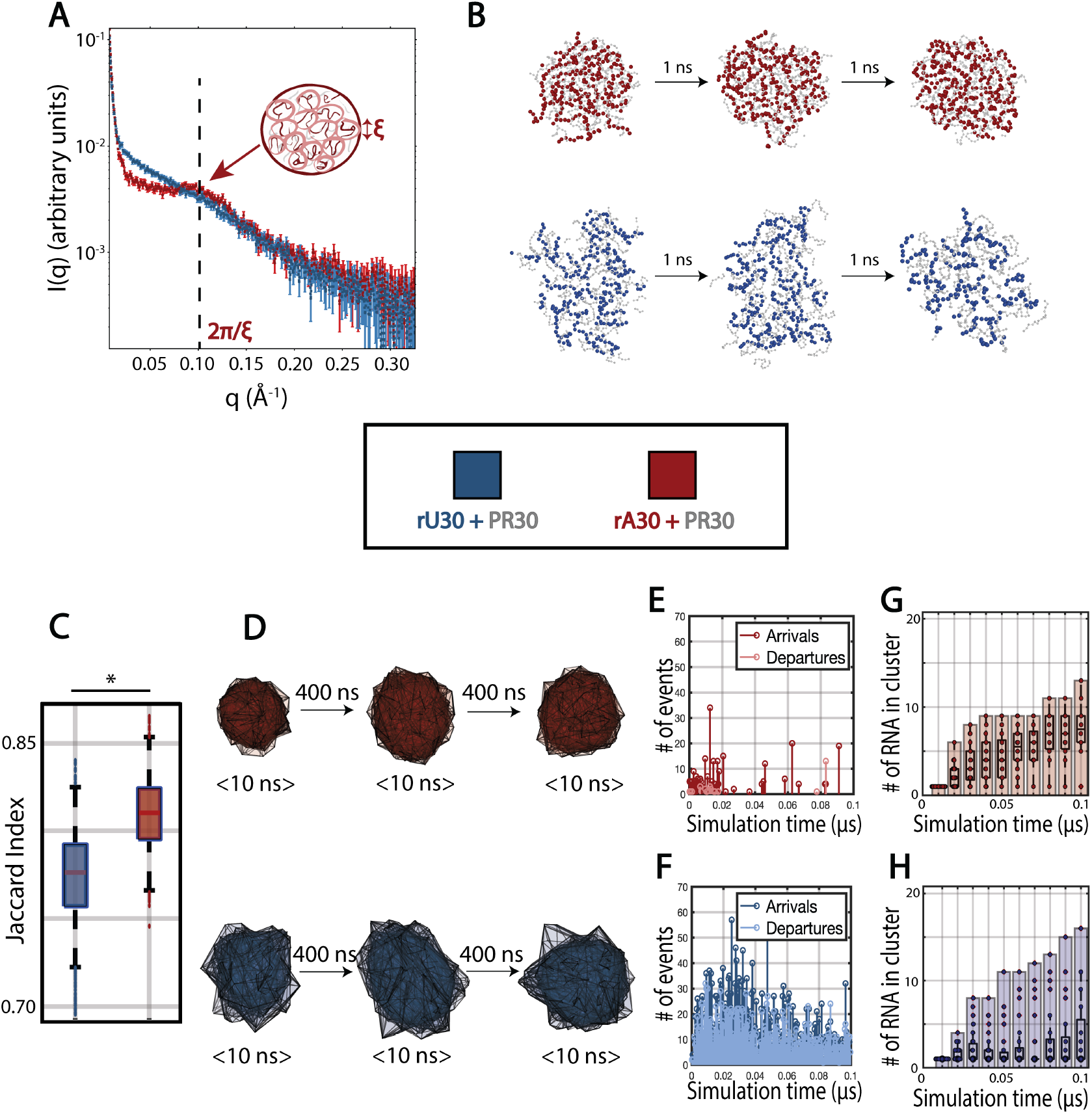
rU30 vs rA30 dynamics within condensates with PR30 at 150 mM NaCl. A) CV-SAXS measurement of rU30 (blue) and rA30 (red) with PR30 at 150 mM NaCl. I(q) is shown on a logarithmic scale, and errorbars represent SAXS measurement errors. The arrow points to a diffuse peak in the SAXS profile which is more pronounced in rA30-PR30. B) Coarse grained molecular dynamics simulations of rU30 and rA30 with PR30 (shown in light grey). For each example condensates of comparable size are shown across three snapshots spaced 1 ns apart, without moving the center of mass in each frame. C) Jaccard indices of rU30 vs rA30 condensate convex hulls across the simulations, quantifying shape similarities. ‘*’ denotes p < 0.001. D) Visualization of condensate convex hulls across the simulation, early (frame 1000-1100), midway (frame 5000-5100) and late (frame 9000-9100) in the simulation. At each timepoint, the same condensate is sampled across ten frames and overlayed on top of each other. E-F) Number of RNA arrivals and departures to and from clusters across the first 0.1 *µ*s of the simulations when condensates are forming. G-H) Number of rU30 (top) and rA30 (bottom) RNA in each cluster of the simulation during the first 0.1 *µ*s of the simulations, when condensates are forming.

The y-intercept of the scattering profile, also known as the zero intensity or I(0) value, scales with molecular weight and can be used to estimate the sample’s association state. A linear extrapolation of the low q scattering to q = 0*Å*^−1^ yields an I(0) value for the rA30-PR30 coacervate samples that is 17.4x larger than rA30-alone, while rU30-PR30 scattering presents a 13.1x I(0) fold increase and rC30-PR30 scattering a 6.8x I(0) fold change (Figure S13). These enhancements signal a large degree of RNA association. The largest change is in the rA30 containing sample, consistent with the more vigorous phase separation seen in poly-A.

At higher q values, the scattering profile of rA30 multiphase samples exhibits a distinctive peak at q ~ 0.1 *Å*^−1^, which is not as pronounced in rU30-containing samples (Figure 4A) or rC30-containing samples (Figure S14A). Such peaks occur in samples that are dynamic but have some degree of ordering, such as amorphous solids or semi-dilute polyelectrolyte solutions. ^61^ They have been observed in phase separated protein mixtures, particularly those formed through coacervation, ^62–65^ and signal an emergent correlation length in the sample. ^33^ We can estimate the approximate lengthscale that corresponds to a peak at this q value. Using equation S.10, we obtain a real space lengthscale of ~ 62.8 *Å*.

### RNA nucleotide sticker strength dictates condensate dynamics

To explain the sequence dependent difference in RNA ordering during phase separation, as reported by CV-SAXS and described above, we sought to model the condensates in real space and to characterize their dynamics. Given our observations that rA condensates are more resistant to salt degradation than rU and rC condensates (Figure 1B, S7A), as well as rA’s high propensity for base stacking interactions, we utilized the Mpipi coarse-grained MD simulation model, whose pair potentials consider such interactions.^41^ Notably, the sticker strengths prescribed for different ribonucleotides in this model empirically agree with our experimental observations, with rA30 having greater *π* − *π* stacking potential than rU30 and rC30.

We performed simulations of each RNA polymer with the PR30 polymer using pair potentials specified by the Mpipi model, ^41^ specified in more detail in Methods section **Molecular dynamics simulations**. Nanosecond scale snapshots of an individual condensate in each simulation are shown in Figure 4B and S14B and full simulations may be viewed in Supplemental Movie 1,2. Qualitatively, condensates with rU30 and rC30 exhibit greater spatial fluctuations (Figure 4C, S14C) than those containing rA30. Due to the weaker interaction potential of pyrimidine bases, rU30 and rC30 molecules separate and re-associate within the condensate, deforming its overall shape more dynamically. When condensates dissociate, a single cluster becomes two clusters, resulting in fluctuation in the number of clusters throughout time (Figure S15). In contrast, rA30 containing condensates stay associated once paired; condensates maintain stable spherical geometry over time (Figure 4D) and the number of clusters remains stable (Figure S15).

To more quantitatively support this point, we follow the changing geometry of a number of individual condensates, by sampling their shapes (parameterized as convex hulls) across simulation frames and computing Jaccard indices between them (Figure S16A, Supplemental Methods section **Condensate shape fluctuation**). The rU30 containing condensates have significantly lower Jaccard indices, which suggest that their shapes are more dissimilar across time (Figure 4C,D, Figure S16B,C). In addition, rU30 and rC30 molecules move in and out of condensates more readily than rA30, with 8070 RNA arrivals into condensates and 7020 departures from condensates for rU30 throughout the 1*µ*s simulation, 3462 arrivals and 2398 departures for rC30, and 371 arrivals and less than 34 departures for rA30. (Figure 4E,F, Figure S17) rA30 transitions occur mainly at the beginning of the simulation when clusters begin to form - once an rA30 cluster is formed, RNA rarely leaves it. As clusters form in rA30, a monotonic increase in the number of RNA in each cluster is seen (Figure 4G, Figure S18). Conversely, growth in rU30 condensates is not monotonic; a proportion of clusters maintain low amounts of RNA (Figure 4H, Figure S18). Similar behavior is seen with rC30, where a population of low-RNA containing clusters persists across the simulation and RNA arrivals and departures occur throughout (Figure S14D,E, Figure S15-S17).

Within condensates and RNA-PR30 complexes, and on average, individual RNA chains undergo a steady increase in *R*_*g*_ and become more extended as the interaction with PR30 progresses and condensates grow (Figure S19A-C). At their start, condensates containing rU30 chains start at *R*_*g*_ = 20.00 ± 0.36 *Å, R*_*EE*_ = 46.84 ± 1.92 *Å* and increase across 200 ns to *R*_*g*_ = 21.08 ± 0.38 *Å, R*_*EE*_ = 51.00 ± 1.85 *Å* in the presence of PR30. Condensates containing rA30 chains begin at *R*_*g*_ = 17.42 ± 0.28 *Å, R*_*EE*_ = 36.58 ± 1.38 *Å* and increase across 200 ns to *R*_*g*_ = 21.12 ± 0.28 *Å, R*_*EE*_ = 49.98 ± 1.54 *Å*. After this 200 ns period, when most RNA has associated into a condensate, the size of RNA chains stabilizes and the late stage *R*_*g*_ and *R*_*EE*_ are comparable to those at 200 ns (Figure S19D). Lastly, the *κ*^2^ value for both poly-U and poly-A undergoes an increase of 16.42% to 0.4488 ± 0.0230 and 58.11% to 0.4408 ± 0.0159 respectively as they interact with PR30, which suggests that the RNA conformations extend as condensates form. With RNA alone in the simulation box, this increase in *R*_*g*_ does not occur, suggesting it is due to protein interaction and not a result of equilibration effects (Figure S20).

## Discussion

The unique information provided by CV-SAXS allows us to observe the structures of RNA within multiphase RNA-protein coacervate mixtures, essentially providing a structural probe of the internal architecture of RNP condensates. We find that poly-U and poly-A have distinct conformational behaviors within RNA condensates. Measurements were made at two different coacervate interaction strengths, achieved by varying the competitive binding by introducing NaCl. This strategy enables a model of RNA condensation with PR30 in two different regimes (Figure 5): at high salt (600 mM NaCl), and at more physiological values (150 mM NaCl). We now place our results into the context of existing theories of coacervation. These theories suggest two major driving forces behind coacervation: interaction of RNA’s negatively charged phosphate backbone with oppositely charged binding partners, here the positively charged amino acid side chains of the partner peptide, as well as base (sequence) dependent *π* interactions with the aromatic nucleotides. Both considerations influence condensate formation and dynamics. A long term goal of this work is to connect RNA conformation in condensates with the distinct sequence-dependent behavior that has been reported in the literature.^22,30^

**Figure 5:**
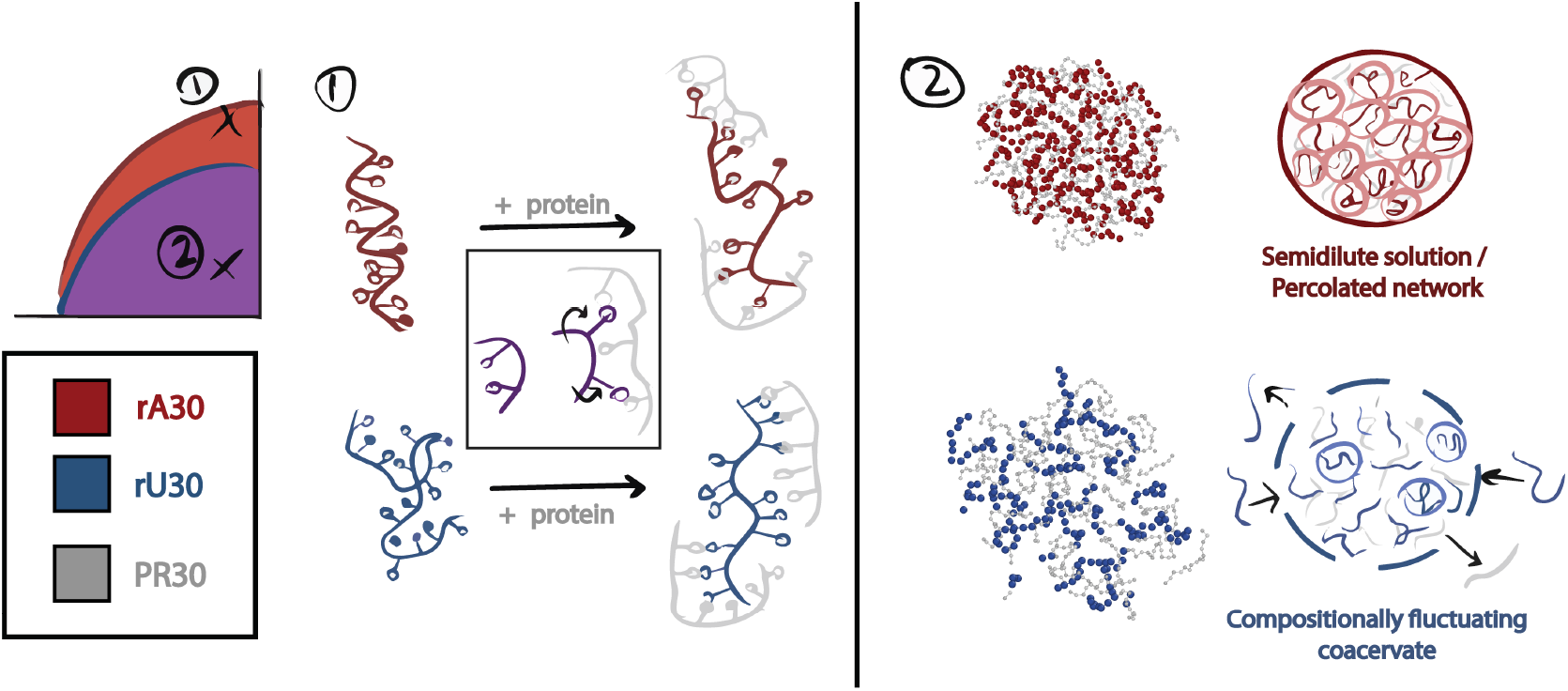
Model of nucleotide-specific RNA conformations and dynamics within ribonucleoprotein condensates. In this study we investigated the greater phase separation propensity of poly-A relative to poly-U from an RNA structure viewpoint. Two regimes were studied - high NaCl near the binodal boundary where subsaturated RNA-protein complexes form and low NaCl where phase separation is favorable. At high NaCl, where individual RNA and protein strands begin to interact, nucleotide bases flip outward along the RNA backbone. rA30 loses its helical conformation while rU30 gains helicity. In both cases, bases flip outward and intramolecular *π* −*π* stacking is abrogated as a result. We infer that this is caused by cation-*π* interactions with R residues that outcompete the native *π* −*π* interactions within RNA. At low NaCl, where RNP condensates form, rA30 condensates are energetically stable with minimal intra-condensate movement while rU30 condensates are compositionally fluctuating. We speculate that this results in droplet-spanning networks with a fixed correlation length *ξ* in rA30-containing condensates, akin to a semidilute solution or coacervate gel. Such correlation lengths are too transient to be reliably detected.

At high salt (600 mM NaCl), when RNA and protein begin to interact, the ensemble modeling of our data suggests that intra-RNA base stacking decreases as nucleotide bases flip outward along the chain. In this regime, rU30 chains adopt helical periodicity while rA30 chains have their periodicity abrogated (Figure 3). Both rU30 and rA30 exhibit subtle increases in *R*_*g*_ upon interaction with PR30 chains (Figure 2B). It has been inferred that base pair driven reptation between RNA chains is marked by increases in *R*_*g*_ and *R*_*EE*_ due to the extension of RNA chains as they twist around each other.^53^ A similar phenomenon may be inferred from our CV-SAXS-based analysis where rU30, having less inherent base stacking, requires less enthalpy to be “molded” by the protein into helical conformations through electrostatic contacts. This interaction could occur if the RNA and peptide strands become intertwined, with heterotypic cation-*π* and *π* −*π* interactions stabilizing their interactions. In contrast, rA30 chains display prominent helicity in the absence of PR30, enabled by robust *π* − *π* stacking of 70.23 ± 2.92 % of its bases (Figure 3). Upon interacting with protein, a similar intertwined conformation may incur greater enthalpic cost associated with flipping its bases outward to interact with a protein binding partner. These higher salt findings provide insight into the RNA-protein interactions which precede phase separation.

At more physiological salt levels, (150 mM NaCl), RNA-PR30 phase separation occurs (Figure S8) for all RNA sequences. The emergence of a low angle peak in the scattering profile of poly-A RNA in the condensate (Figure 4A), suggests a lengthscale characteristic of short range order in semidilute polymer solutions. ^33,64^ This loose ordering suggests more stable structuring in rA30-PR30 condensates than rU30-PR30, and is supported by MD simulations showing that RNA chains within rA30 condensates form a dynamic mesh (Figure 4B). In these simulations, this mesh-like feature is stable across time - once an rA30 RNA strand incorporates with a cluster it rarely leaves (Figure 4E), resulting in consistently spherical cluster geometry (Figure 4D). Conversely, simulations suggest that rU30 and rC30 clusters are dynamic with rapidly fluctuating shapes (Figure 4C). As a result, components of the condensate can dissociate (Supplemental Movie 1) and RNA more freely enters and exits this phase (Figure 4F). These greater dynamics oppose prolonged short range ordering, consistent with the lack of a peak in their SAXS profiles (Figure S7A). For these reasons, rU and rC condensates appear to be fluctuating in shape and composition as they form. We infer this from the fact that rU and rC condensate geometries are more unstable over time, brought on by consistent RNA departures that result in more dynamic clusters with fluctuating shape and non-monotonic growth in the number of RNA per cluster (Figure 4H). Conversely, the stronger rA interactions produce condensates resembling long-lived droplet-spanning networks, similar to percolated clusters (Figure 5). ^9^ Indeed mesh-like RNA condensates have been observed, both dynamic and gel-like, ^66^ and purine-rich RNA has been linked to slower condensate dynamics and greater persistence when coacervation driving forces are lowered with increasing NaCl. ^30,31^ Finally, these simulations provide additional insight into experimental reports that poly-A condensates form more readily than poly-U and poly-C condensates and may have different architectures (Figure 1).^30^

Unlike the dilute phase with little association between RNAs, our analysis cannot resolve all-atom chain conformations in the dense phase. We instead report trends in molecule size and overall condensate dynamics, with the aid of our MD simulations. The literature reports different trends where macromolecules are purported to increase in size relative to the dilute phase in some cases^53,67^ and decrease in others.^68^ Our higher salt experimental results suggest that RNA extends when it interacts with PR30 (Figure 3). Our simulation results show that RNA undergoes an initial compaction as it is met with a large amount of positive charge in PR30 and then extends as it interacts with PR30 over time, conceivably due to a similar intertwining phenomenon and *R*_*g*_ increase seen at high salt (Figure S19). This phenomenon is more pronounced for poly-A.

Uniform electrostatic interactions between the RNA backbone and positively charged protein residues, which increase linearly with RNA length, constitute much of the driving force for RNA-driven LLPS.^69,70^ Beyond this, our work stresses the role of RNA sequence in fine tuning its attractive potential within condensates and the structural arrangements of RNA. We provide further insight as to why rA30 condensates are more robust to salt denaturation (Figure 1B). Poly-A/G + PR30 condensates were previously shown to be less fluid than poly-U/C + PR30 condensates, supporting this description. ^22^ Through varying its purine/pyrimidine content ratio, RNA can vary its sticker strength within condensates - purine-rich RNA condensates may have greater gelation tendency while pyrimidine-rich RNA condensates may be more liquid-like. This may allow the cell to tune RNP condensate fluidity, such as with poly-A rich RNA localization to stress granules. ^3^ Moreover, the striking difference between rA30 vs rU30 flux in and out of condensates (Figure 4E,F) also alludes to control of condensate composition, stability, and material properties through client exchange vs retention.^71,72^ Such knowledge provides insight on how to encourage or discourage their formation in disease contexts.

Finally, we address the possible role of sucrose in these studies. Such high concentrations of sucrose are required to perform CV-SAXS, which benefit the field by revealing the conformations of RNA alone within the mixture. Generally, CV-SAXS is applicable to systems that undergo strongly associative phase separation, past experimental results show consistency between contrast matched and unmatched conformations. It is important to point out that the chains within larger clusters are highly polydisperse, so the measurements cannot be easily interpreted using scattering analyses that assume a monodisperse sample once association occurs (in contrast to the lower NaCl condition, where RNA protein association dominates over RNA-RNA association). We observe that the time scale for condensate formation increases, possibly due to slower macromolecular motions given the increased viscosity of the solution. However, sucrose does not influence the overall morphology of droplets, which remain liquid-like (Figure S8), suggesting that the driving forces for phase separation are not affected. Under the conditions studied here, sucrose does not accelerate phase separation compared to samples with no sucrose so does it not act as an artificial crowding agent.

Additional studies using all-atom MD simulations would be valuable in further characterizing the interactions between RNA and proteins. Having demonstrated that CV-SAXS measurements are amenable to examining RNA within condensates, we look toward the potential applications of this method to RNA-protein systems such as those involved in neurodegeneration and those involving longer noncoding and mRNA constructs.^12,73,74^ As well, SAXS methods may prove useful in experimentally validating simulations of RNA and protein conformations in the dense vs dilute phases.^53,67,75^

## Conclusion

Given the importance of ribonucleoprotein condensates, knowledge about the internal structure of RNA constituents is desired. We demonstrated the utility of contrast-variation SAXS in observing RNA conformational changes and condensate properties upon phase separation with polybasic PR30. Utilizing this methodology we characterized distinct behaviors in poly-U and poly-A RNA conformational changes preceding phase separation, where RNA chains extend and nucleotides flip outward to facilitate heterotypic interactions with PR30. In the condensed phase, poly-A RNA condensates are more salt resistant, more temporally stable, and more spatially ordered than poly-U condensates. Beyond simply considering uniform electrostatic interactions of the negatively charged backbone, the nucleotide sequence and chain conformations of RNA may modulate its condensate behavior. The implications of backbone length and nucleotide sequence on coacervation constitutes the molecular grammar of RNA-associated phase separation.^76^

## Author Contributions

LP and TW conceived this study. TW performed biochemical and microscopy assays on RNA condensates. QH and TW prepared samples for CV-SAXS. TW, SF, QH performed and analyzed synchrotron CV-SAXS measurements. TW performed MD simulations, ensemble optimization, and computational analysis. LP acquired funding and provided supervision. TW and LP wrote the manuscript. All authors were involved in editing the manuscript.

## Acknowledgements

We are grateful to Qingqiu Huang and Samantha MacMillan for support with SAXS data collection; Rachael Skye, George Padilla, and Gundeep Singh for valuable advice on MD simulations; Greg Powers for support with contrast-variation SAXS theory; Dhruva Ajit Nair and Robert Dunleavy for assistance with the SpectraMax plate reader; and Charlotta Lorenz, Teagan Bate, Sully Bailey-Darland, Takumi Matsuzawa, Kaarthik Varma, Dana Matthias, Jen Jang, and Eric Dufresne for constructive discussions about phase separation.

